# A 3-Dimensional bioprinted human gut-liver axis model for studying Alcoholic Liver Disease

**DOI:** 10.1101/2022.04.26.489511

**Authors:** Kranti Meher, Madhuri Rottela, K Saranya, Subramanian Iyer, Gopi Kadiyala, Subhramanyam Vangala, Satish Chandran, Uday Saxena

## Abstract

Gut-liver axis is the interaction between the gut, its microbiome and the liver. The crosstalk and interaction between these organs plays an important role in their individual health and disease. Alcoholic liver disease (ALD) is a case in point where dysfunction of intestine actively promotes liver damage by alcohol. A flashpoint in ALD is the breach of intestinal integrity caused by gut bacteria ***Enterococcus faecalis*** (*E.Faecalis*). More specifically, Cytolysin, a toxin secreted by this bacteria may have a central role in the genesis of ALD. 3-D bioprinted human simulations of the gut-liver axis may help better understand the genesis of ALD.

Here we developed a 3 dimensional bioprinted in vitro model composed of human origin intestinal and liver cells to explore the role of Cytolysin and ethanol in intestinal and liver damage. We find that neither Cytolysin or ethanol are sufficient for cell damage but a combination of the two act in concert to cause maximum breach in intestinal integrity. Secondly we find that enhanced transport of macromolecules thru the intestinal layer is not caused by overt cell toxicity but occurs through potentially paracellular/transcellular pathways. Our model will be used to test repurposed and new drugs/ biologics for treatment of ALD as well as other intestinal inflammatory diseases.

## Introduction

Alcohol liver disease (ALD) is a prevalent liver disease globally and there are no targeted therapies currently on the market for it. While there are animal models for this disease which include binge ethanol feeding, there is a need for a specific human like model to be able to study the various steps in disease genesis and the role of various molecular pathways in the disease process.

In ALD the gut bacteria *E.Faecalis* is believed to secrete a toxic protein Cytolysin and together with ethanol they damage the intestinal and liver cells. Its hypothesized that Cytolysin reaches the liver and is able to inflict liver injury as part of the ALD process. The exact mechanisms of how Cytolysin may reach the liver is not clear and may involve breaching of intestinal integrity by severely damaging the intestine or by paracellular or transcellular pathways. In the studies reported here we have explored the impact of Cytolysin and ethanol on trans-intestinal transport pathways and the cell damage to intestine and liver cells caused by the agents.

## Brief Methods

3D bioprinting of the cells with bioink was performed as described before. Transwell filters (Sigma Aldrich) housed the human intestinal Cac02 cells and the human liver HepG2 cells were printed on the plate directly.

Transport studies using FITC Dextran were performed as described below. MTT assays to asses cell viability and alkaline phosphatase assays were performed using a commercial kit from Sigma Aldrich.

## Results

### 1. Establishing the 3 D Bioprinted gut-liver axis model

To establish the model we used a modular approach. Human Intestine derived Caco2 cells were bioprinted in a transwell filter on bioink and inserted into 24 well plate. A second module consisted of human liver HepG2 cells printed on the wells of the plate directly on bioink. Figure 1 shows the arrangement of our 3D bioprinted model.

**Figure 1:**
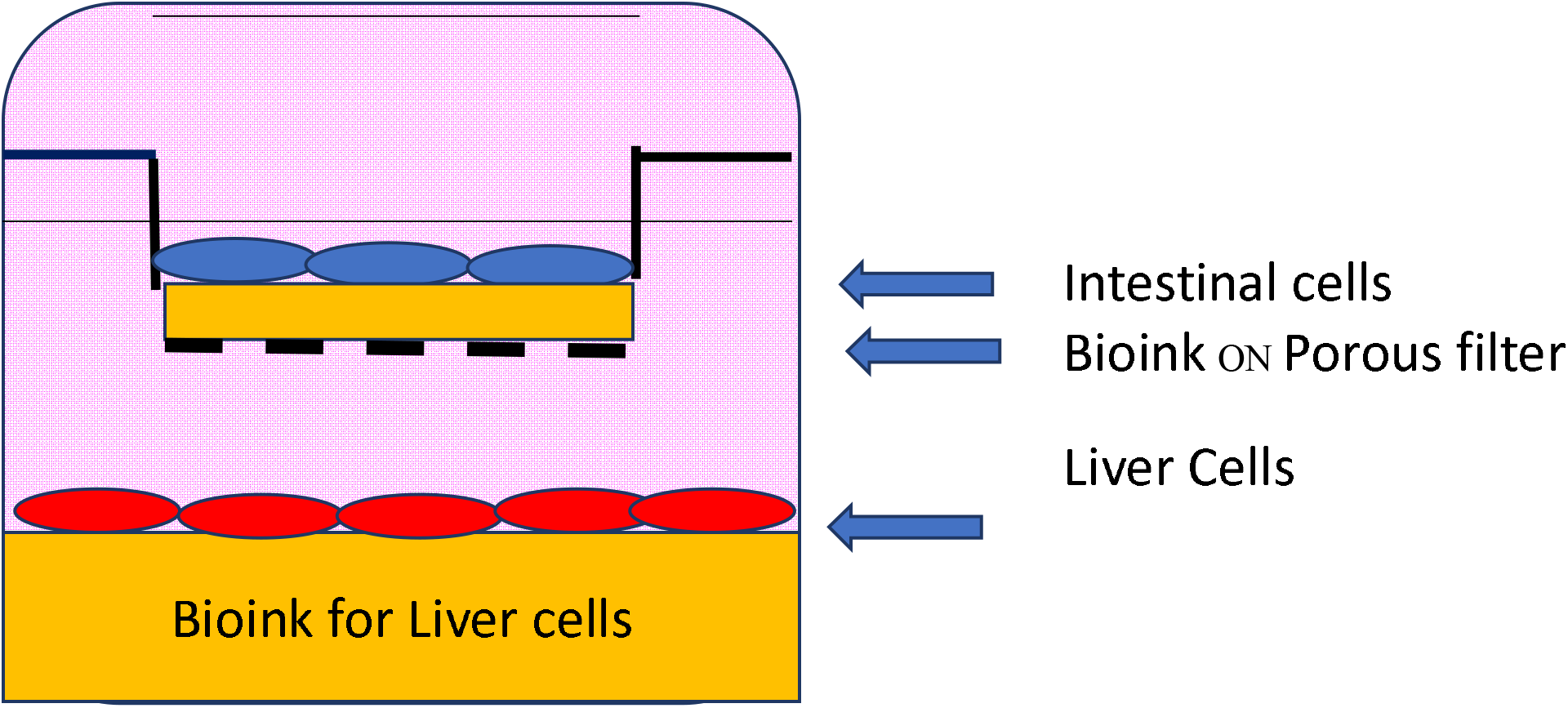
GUT-LIVER AXIS HUMAN CELL BIOPRINTED MODEL

This two module approach allows us to separately examine each of the cell types as needed by simply removing the transwell filter with Caco2 cells on it and studying it and then directly also examining the lower layer of HepG2 liver cells. We can also sample media from the intestinal apical side top module and separately from the bottom liver cells module. The porous transwell filter allows free flow of molecules between the two compartments, much like the portal vein that connects the intestine to the liver.

To establish the barrier function of our model, we used 4 kDa fluorescein isothiocyanate (FITC)-labelled dextran (FD4, Sigma Aldrich). FITC Dextran was added to the top half (intestinal apical side) and its presence in the bottom half was sampled over a period of time. Fluorescence intensity (excitation 492 nm/emission 520 nm) was measured using a fluorimeter. If the intestinal layer is not breached, very little of dextran should be found in media in the bottom module.

Breach of intestinal integrity is the starting point of several disease, essentially allowing toxins and other macromolecules to get in systemic circulation and reach other organs. To test the integrity of our Caco-2 confluent bio printed monolayer, which mimics the intestinal barrier, we measured the trans-epithelial transport of (FITC)-labelled dextran (FD4, Average MW 4000). Briefly fluorescent FD4 (0.01 or 0.2 mg/ml in media) was added to the top (apical) side of intestinal module. Then, to estimate trans epithelial transport, media was sampled from the lower module at time points indicated and florescence read.

As shown in Figure 2, the following conditions were used

- Control wells with no intestinal cells printed on the filter but dextran added to measure transport
- Control cells with no dextran added to measure background florescence
- Wells with cells printed on the filter and dextran added

**Figure 2:**
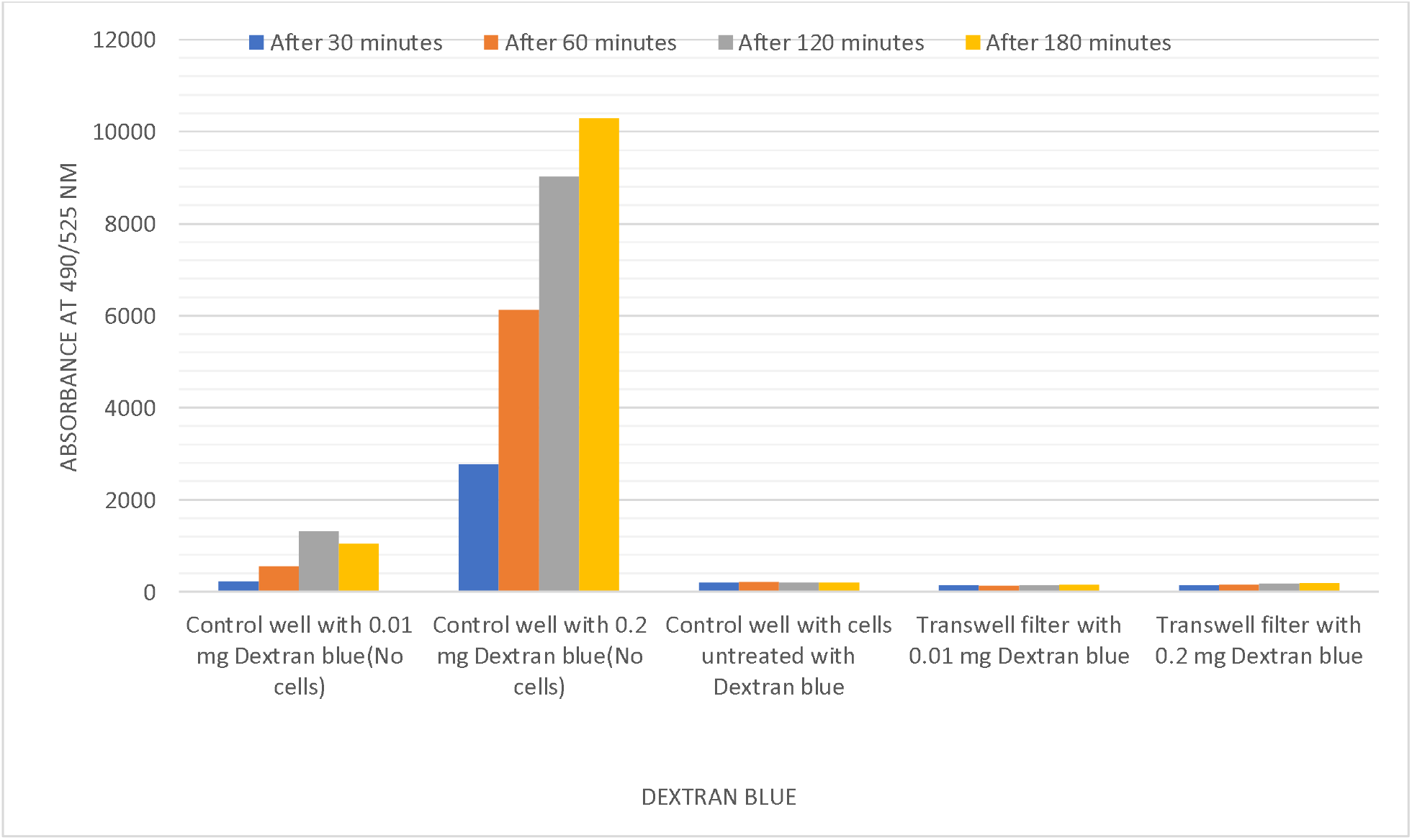
TRANSPORT OF FITC DEXTRAN ACROSS INTESTINAL CELL LAYER

Shown above is the transport across the 3D model at different time points after addition of dextran. The transport across was highest in the filters that contained no cells printed as expected (First set of bars 0.01 mg/ml dextran added) and second set of bars (0.2 mg/ml dextran added) from the left in the figure above with commensurate increase with higher concentration of dextran. The third set of bars represents background florescence and was very low.

The addition of dextran to filters on which intestinal cells were printed, there was hardly any florescence observed in the lower module (fourth and fifth set of bars, Figure 2) suggesting that the printed intestinal cells act as a robust barrier to transport and present a tight integrity intestinal layer.

### 2. Examining the effect of Cytolysin and Ethanol on trans intestinal transport

During genesis of ALD, the *E. Faecalis* gut bacteria secrete Cytolysin, a toxic protein that can potentially damage and induce cell death. We used the 3D bioprinted model validated for intestinal integrity as described above to study the role of purified Cytolysin, ethanol (1%) alone or in combination. As shown below in Figure 3, Cytolysin (100 ug/ml) or ethanol alone or a mixture of the two were incubated with the cells for 24 hours and then transport studies with dextran were performed. Figure 3 shows that neither Cytolysin or ethanol (1%) alone were able to enhance dextran transport across the intestinal cell layer relative to untreated cells (shown as control transwell filter) at every time point sampled from 30-180 minutes. In contrast the combination of the two dramatically increased the transport by about 200 percent at every time point. This suggest that Cytolysin may work together with ethanol in ALD to enhance transport across intestine.

**Figure 3:**
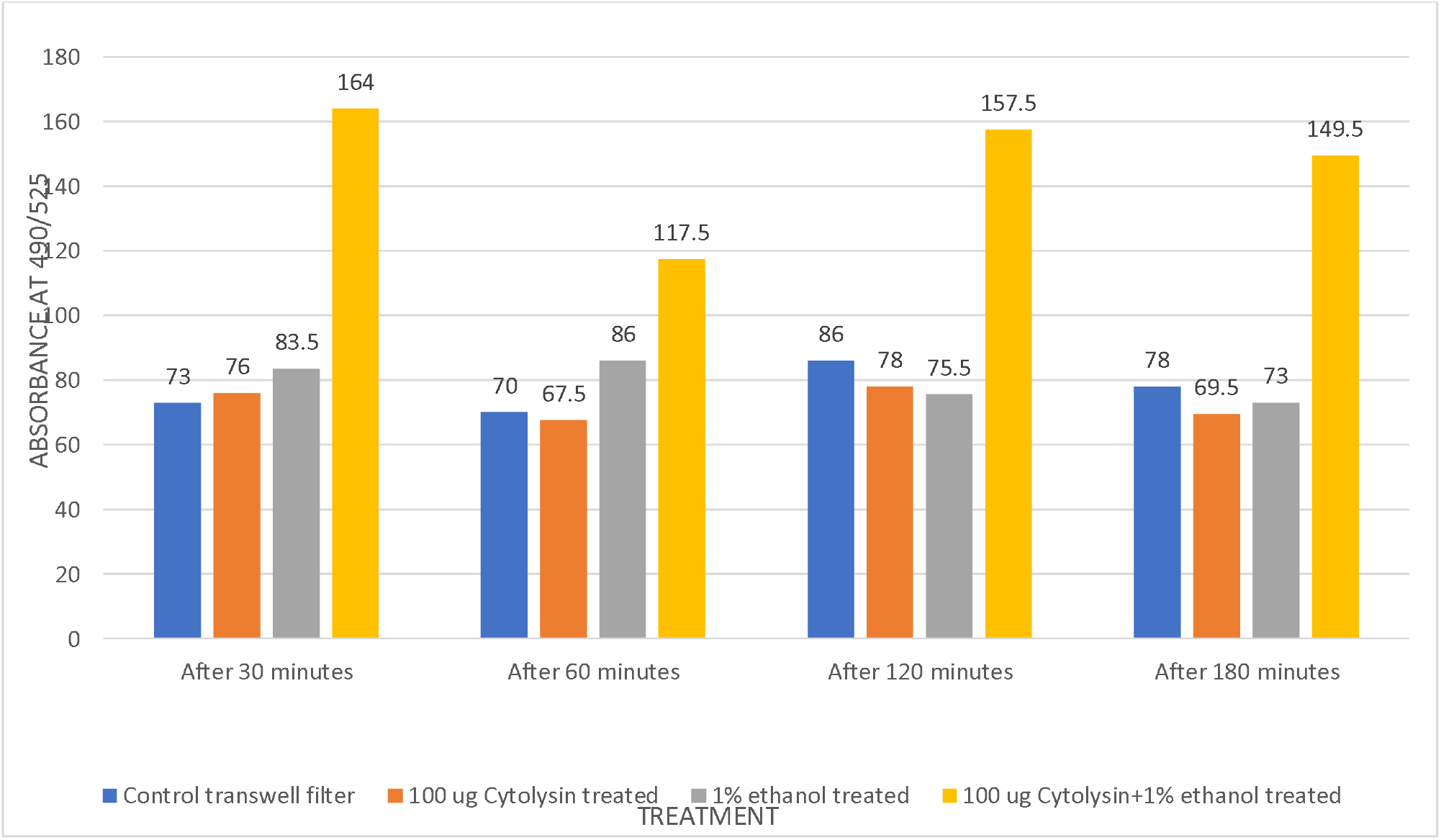
TRANSPORT OF FITC-DEXTRAN ACROSS INTESTINAL CELL LAYER

### 3. Increased transport by Cytolysis and ethanol does not require cell death

The increased transport caused by the combination of Cytolysin and ethanol could be due to a) direct cell damage/death of the intestinal cell monolayer leading to loss of monolayer integrity or 2) due to transcellular or paracellular pathways in living cells induced by the combination.

To explore if there was any cell death caused by the combination, we performed an MTT assay which asses cell viability of both the intestinal cells and the liver cells. As shown in Figure 4, there was no decrease in cell viability of intestinal cells (higher absorbance is a result of higher dye uptake by living cell and increased colour formation in MTT assay). Similarly there was no change in liver cell viability by MTT assay.

**Figure 4:**
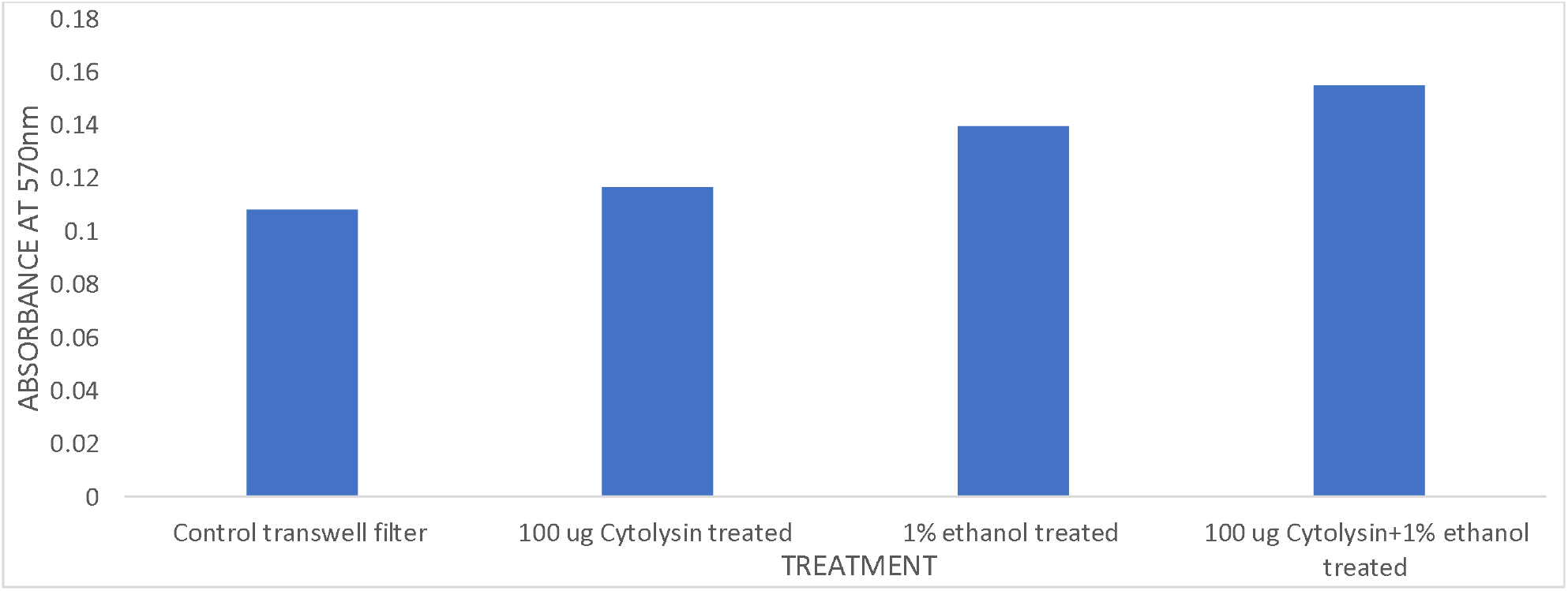
ASSESMENT OF CELL VIABILITY BY MTT OF INTESTINAL CELLS

**Figure 5:**
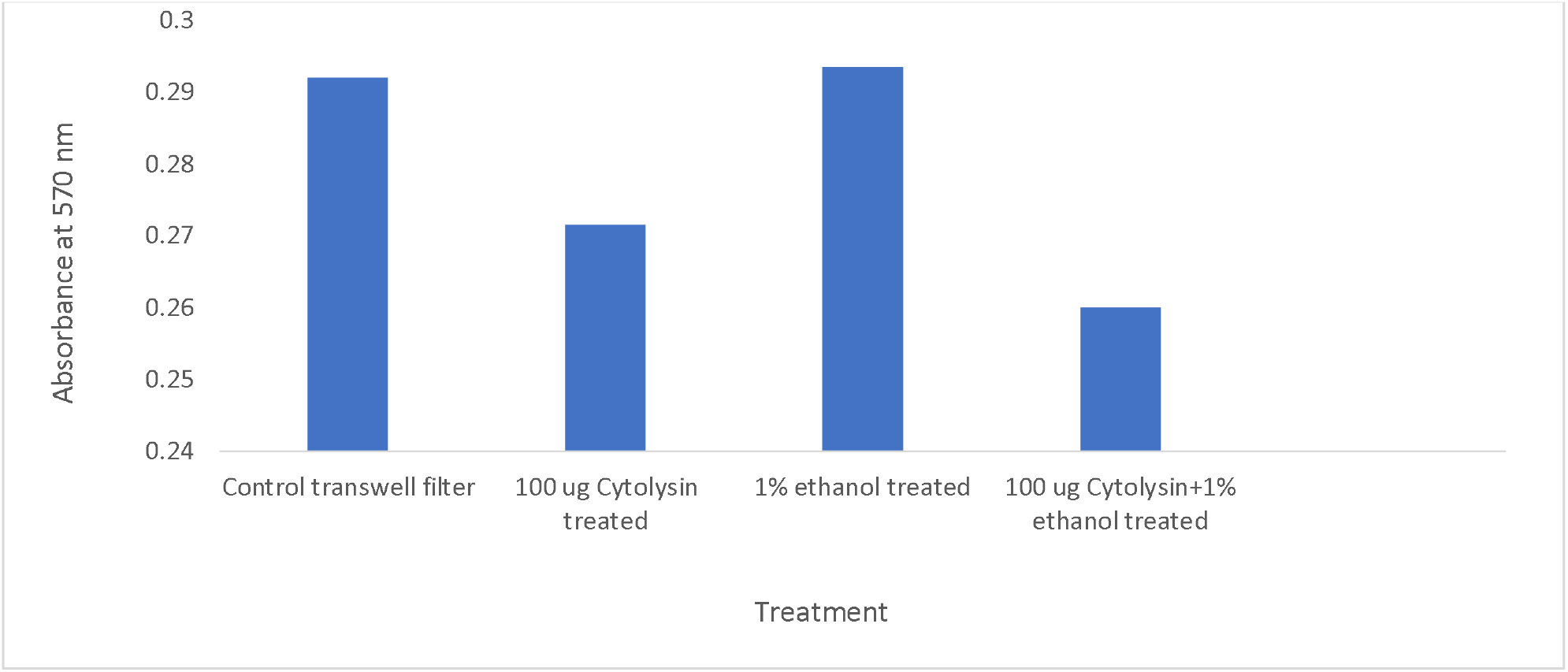
ASSESMENT OF CELL VIABILITY BY MTT OF LIVER CELLS IN LOWER MODULE

These data support the thesis that Cytolysin and ethanol rather than causing gross cell death cause subtle changes in the intestinal cells to promote increased transcellular or paracellular pathways of transport.

### 4. Higher concentration of Cytolysin also does not cause cell damage as assessed by liver cell biomarker

The ultimate target organ in ALD is the liver. It is possible that Cytolysin breaches the intestinal layer and reaches the liver and together with ethanol to cause liver cell death. We therefore looked to asses if higher amounts of Cytolysin may cause liver cell death. We examined the effect of 100 ug/ ml and a higher concentration of 500 ug/ml of Cytolysin with and without ethanol on dextran transport as well as secreted alkaline phosphatase, a biomarker often used to asses liver cell damage/death. As shown in Figure 6, Cytolysin alone at 500 ug/ml did not significantly increase transport of dextran but there was an increase when added together with ethanol (compare transport in control cells versus Cytolysin at 500 ug/ml versus 500 ug/ml plus ethanol at 120 minutes, grey bars in figure 6).

**Figure 6:**
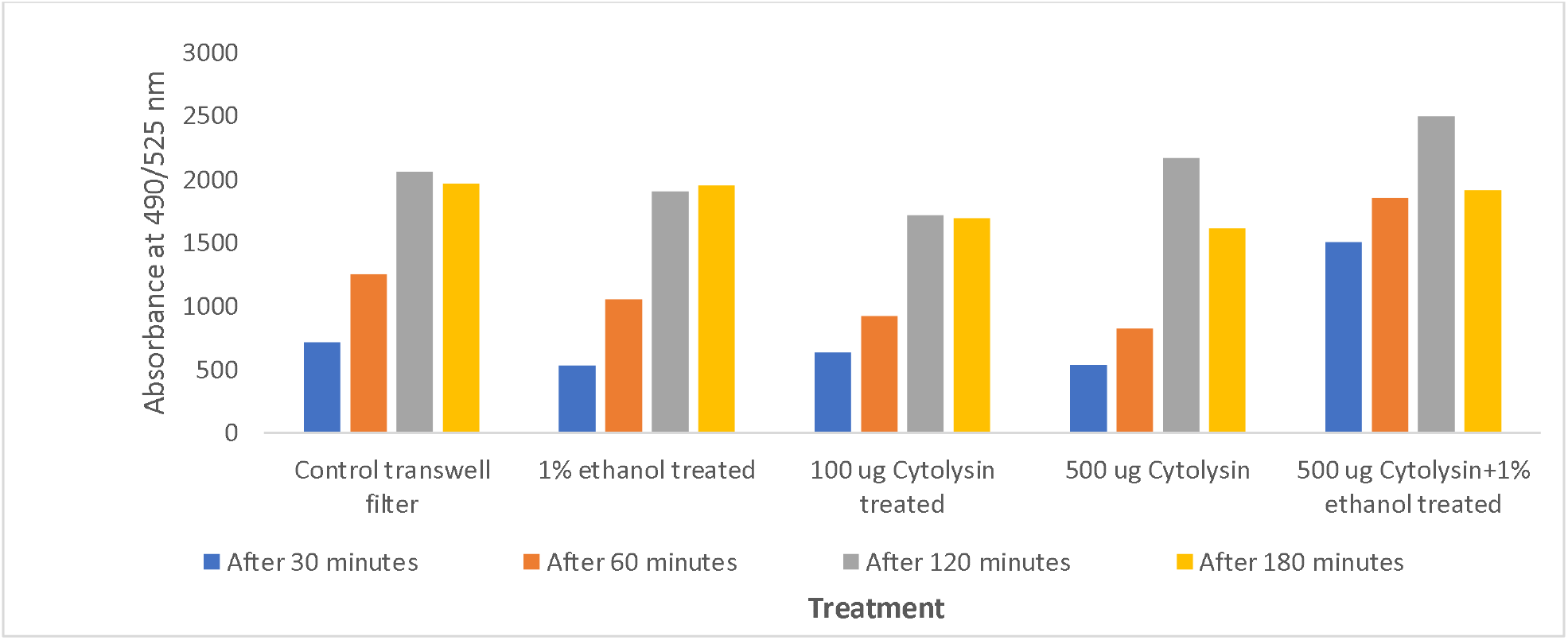
TRANSPORT OF FITC DEXTRAN

In the same experiment to detect any liver cell damage we used alkaline phosphatase secretion as a marker for liver damage. Alkaline phosphatase enzyme is a sensitive marker whose secretion is considered to reflect liver cell damage. As shown in Figure 7, the first 4 bars show standard curve of alkaline phosphatase activity to demonstrate that even low levels of enzyme activity can be detected. The next 5 bars show activity of alkaline phosphatase in the cell culture media of from lower module which houses the liver cells. There was no increase of the biomarker’s activity again reiterating that the damage to liver cells by Cytolysin and ethanol under conditions used is minimal.

**Figure 7:**
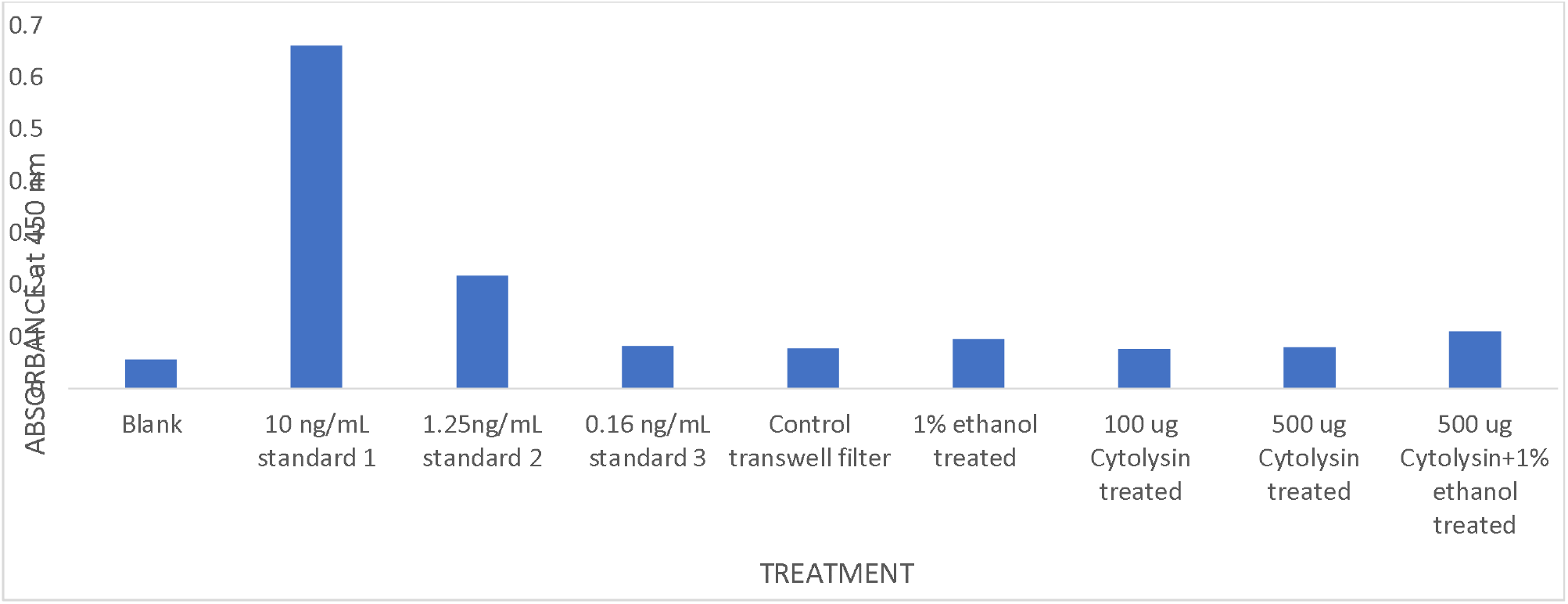
ALKALINE PHOSPHATASE ACTIVITY IN MEDIA FROM LOWER LIVER CELL MODULE

## Discussion

Here we describe for the first time gut/liver axis model using a 3D bioprinted human cells to tease out roles of Cytolysin and ethanol, the dominant insults in ALD. Our results have uncovered some exciting new findings

1. Unexpectedly, we find that Cytolysis alone or ethanol alone do not change the trans intestinal transport of dextran across intact intestinal monolayer but only the combination of two induces significant increase in transport indicating breach of intestinal integrity
2. Our data further suggest that the enhanced transport by the Cytolysin and ethanol combination does not induce overt cell death suggesting that transport may be occurring thru paracellular (loosening of tight junctions between intestinal cells) and or transcellular pathways

Our data are similar to in vivo data recently reported in a model of ALD. In a mice study for Cytolysin and ethanol contribution to liver damage, Duan et al examined effects of the cytolytic *E. faecalis* versus contribution by ethanol. In their studies, the highest degree of liver injury and inflammation as well animal death were seen only when ethanol was dosed together with the cytolytic *E. faecalis* relative to the bacteria alone treated or ethanol-alone controls animals. This suggests that these two interventions work together to result in maximum damage to intestine and liver.

One caveat to our findings is that there is hardly any cell death induced by Cytolysin and ethanol – but the conditions used were more acute in nature (overnight incubation of Cytolysin and ethanol) whereas in disease conditions the intestine and liver may be exposed to chronic onslaught of these insults and cause significant cell damage.

The model we have created of human gut/liver axis can not only be used ALD but for several other diseases such as NAFLD (non-alcoholic fatty liver disease) as well as diet induced fatty liver. The model can also be used to study intestinal inflammatory diseases such as IBD, Crohns and ulcerative colitis.

